## Supplementary Methods, Figures, and Video Caption for "A geometric attractor mechanism for self-organization of entorhinal grid modules"

(Dated: March 8, 2019)

### CONTENTS

Simulation setup 1
- Standard model 1
- Simplified model for phase diagrams 2

Simulation data analysis 2
- Spatial rate maps and autocorrelation functions 2
- Grid scale, orientation, and gridness 2
- Module clustering 3

Varying velocity gain model 4
- Simulation setup with a velocity gain gradient 4
- Results with a velocity gain gradient 4

Supplementary Video caption 5

Supplementary Figures 7

References 15

### SIMULATION SETUP

#### Standard model

To distribute neural subpopulations evenly, we assign each position in a $2 \times 2$ block of neurons to a different subpopulation and tile each network with these blocks. In other words, for a network of size $n \times n$, the preferred network directions are $\hat{\mathbf{e}}(2i-1, 2j-1) = -\hat{\mathbf{x}}$, $\hat{\mathbf{e}}(2i-1, 2j) = \hat{\mathbf{y}}$, $\hat{\mathbf{e}}(2i, 2j-1) = -\hat{\mathbf{y}}$, and $\hat{\mathbf{e}}(2i, 2j) = \hat{\mathbf{x}}$ for block indices $i, j = 1, \ldots, n/2$. The preferred spatial directions take corresponding values $\hat{\mathbf{E}}(2i-1, 2j-1) = -\hat{\mathbf{X}}$, $\hat{\mathbf{E}}(2i-1, 2j) = \hat{\mathbf{Y}}$, $\hat{\mathbf{E}}(2i, 2j-1) = -\hat{\mathbf{Y}}$, and $\hat{\mathbf{E}}(2i, 2j) = \hat{\mathbf{X}}$.

We initialize each neuron with a uniformly-distributed random firing rate between 0 and 0.001 (arbitrary units). We evolve 500 timesteps without velocity input to generate grid-like activity. Next, we anneal grid defects. For each velocity angle $\pi/2 - \pi/5$, $2\pi/5$, and $\pi/4$, we evolve 5000–10000 timesteps with constant speed $0.5\,\mathrm{m/s}$. We then evolve 50 000 timesteps with velocity data from a real rat trajectory within a circular enclosure [1, 2]. The main simulation phase ensues with continuation of velocity input from the trajectory. For each network $z$, we randomly choose three neurons within a distance of $0.15n$ from the network center. Throughout the main phase, we tabulate their mean firing rates as a function of rat spatial position.

#### Simplified model for phase diagrams

To generate data for the phase diagrams in Fig. 4 of the main text, we set up our simulations in a similar way, with the following differences. We use only 2 network depths. We use slightly larger velocity gain $\alpha = 0.4\,\mathrm{s/m}$ to produce grids of smaller spatial scale since a greater number of activity peaks allows for better measurement of grid scale. After initializing the system and performing initial time evolution in the same manner as in the standard model, we take the activity patterns of the two networks. There is no main phase with extended rat trajectories and single neuron recordings.

### SIMULATION DATA ANALYSIS

#### Spatial rate maps and autocorrelation functions

We discretize the animal's environment into $1\,\mathrm{cm} \times 1\,\mathrm{cm}$ position bins indexed by $\mathbf{R} = (X, Y)$. By tabulating a single neuron's average firing rate when the animal occupies each position, we produce the spatial rate map $S(\mathbf{R})$. We define its normalized spatial autocorrelation function as

$$C(\mathbf{R}) = \frac{\dfrac{1}{N(\mathbf{R})} \displaystyle\sum_{\mathbf{R}'} S(\mathbf{R}')S(\mathbf{R}' - \mathbf{R})}{\dfrac{1}{N(0)} \displaystyle\sum_{\mathbf{R}'} S(\mathbf{R}')S(\mathbf{R}')}, \tag{1}$$

where $N(\mathbf{R})$ is the number of pairs of positions separated by $\mathbf{R}$. $C(\mathbf{R})$ can be efficiently calculated via discrete Fourier transforms.

We can define similar network autocorrelation functions $c(\mathbf{r})$ for the population activity within the neural sheet of each networks indexed $z$.

#### Grid scale, orientation, and gridness

We use autocorrelation functions to extract the scale, orientation, and gridness of spatial and network grids. We first convert each position $\mathbf{R}$ to polar coordinates and calculate the autocorrelation as a function of radial distance $R$ by averaging over polar angle $\Phi$:

$$C_{\mathrm{rad}}(R) = \frac{1}{N(R)} \sum_{\Phi} C(R, \Phi), \tag{2}$$

where $N(R)$ is the number of positions corresponding to each discretized $R$. This function is analogous to the radial distribution function of condensed matter physics. To filter out small fluctuations at the centimeter scale while permitting estimation of the location of extrema at the subcentimeter scale, we use coarse $1\,\mathrm{cm}$ bins for $C_{\mathrm{rad}}(R)$, linearly interpolate its value at every $0.1\,\mathrm{cm}$, and apply a Gaussian filter with respect to $R$ with standard deviation $8\,\mathrm{cm}$. We define the spatial grid scale $\Lambda$ as the $R$ corresponding to the first maximum of the smoothed $C_{\mathrm{rad}}(R)$, not including the maximum at $R = 0$.

Grid orientation and gridness are computed from the angular structure of $C(R, \Phi)$ in the region around $R = \Lambda$. This region is an annulus bounded by $R$'s corresponding to the first and second minima of the smoothed $C_{\text{rad}}(R)$, which we call $R_1^*$ and $R_2^*$. This annulus is analogous to the first coordination shell of condensed matter physics. We average over $R$ within the annulus to calculate the autocorrelation as a function of $\Phi$:

$$C_{\text{pol}}(\Phi) = \frac{1}{N(\Phi)} \sum_{R_1^* \leq R \leq R_2^*} C(R, \Phi), \tag{3}$$

where $N(\Phi)$ is the number of positions within the annulus corresponding to each discretized $\Phi$. To assess the degree of 6-fold symmetry, we calculate the sixth component of the discrete Fourier transform of $C_{\text{pol}}(\Phi)$ using 5° bins for $\Phi$:

$$\Psi_6 = \sum_{\Phi} C_{\text{pol}}(\Phi) \mathrm{e}^{-6\mathrm{i}\Phi}. \tag{4}$$

Orientation angle $\Theta$ is defined as the complex argument of $\Psi_6$ divided by 6. Gridness is the fraction of $C_{\text{pol}}(\Phi)$'s total Fourier power, after removing the zeroth component that describes its constant amplitude, belonging to the sixth component. It is thus

$$\text{gridness} = \frac{2|\Psi_6|^2}{\displaystyle\sum_{\Phi} C_{\text{pol}}(\Phi)^2 - \frac{1}{N_{\text{pol}}} \left[ \sum_{\Phi} C_{\text{pol}}(\Phi) \right]^2}, \tag{5}$$

where $N_{\text{pol}} = 72$ is the number $\Phi$ bins. We need the factor of 2 to account for negative Fourier components which have power equal to that of positive components. By properties of Fourier transforms, $\sum_{\Phi} C_{\text{pol}}(\Phi)^2$ is the total Fourier power, and $[\sum_{\Phi} C_{\text{pol}}(\Phi)]^2 / N_{\text{pol}}$ is the power of the zeroth component. A similar definition for gridness has been proposed to assign a local grid score to each spike [3]. We use this definition instead of others used in the literature [4] because it has an intuitive meaning as the fraction of angular power contributed by 6-fold symmetry to the autocorrelation function.

Network grid scales $\lambda$, orientations $\theta$, and gridness can be similarly extracted via the network activity autocorrelation functions $c(\mathbf{r})$.

### Module clustering

Following Ref. 4, we categorize grid cells into modules by clustering their grid scales and orientations using a $k$-means algorithm. The number of clusters $k$ is determined through kernel smoothed densities (KSDs).

We define linearly rescaled grid scales $\tilde{\Lambda}$ such that the largest and smallest scales for each simulation correspond to 0 and 1. We similarly define linearly rescaled grid orientations $\tilde{\Theta}$ such that 0° and 60° correspond to 0 and 1. We divide $\tilde{\Lambda}$-$\tilde{\Theta}$ space into $0.02 \times 0.02$ bins and define the KSD for each bin as

$$\text{KSD}(\tilde{\Lambda}, \tilde{\Theta}) = \frac{1}{N} \sum_i \exp\left[ -\frac{(\tilde{\Lambda} - \tilde{\Lambda}_i)^2}{2\sigma_{\Lambda}^2} \right] \exp\left[ -\frac{||\tilde{\Theta} - \tilde{\Theta}_i||^2}{2\sigma_{\Theta}^2} \right]. \tag{6}$$

$N$ is the number of grid cells, each of which has scale $\tilde{\Lambda}_i$ and $\tilde{\Theta}_i$. For the periodic variable $\tilde{\Theta}$, we define the distance $||c|| = |c|$ for $|c| \leq 0.5$, $1 - |c|$ for $|c| > 0.5$. We take both standard deviations

$\sigma_\Lambda$ and $\sigma_\Theta$ to be 0.1. We use the number of peaks of this KSD as the initial number of clusters $k$ for $k$-means clustering in $\tilde{\Lambda}$-$\tilde{\Theta}$ space.

We perform $k$-means clustering with random initial points in $\tilde{\Lambda}$-$\tilde{\Theta}$ space 200 times per simulation. For each clustering attempt, we calculate the silhouette, a metric describing degree of separation among clusters [4, 5]. For each grid cell $i$ in cluster $b$, we calculate its average distance in $\tilde{\Lambda}$-$\tilde{\Theta}$ space to all grid cells $j$ in cluster $c$:

$$D^c_{bi} = \frac{1}{N_c} \sum_j \sqrt{(\tilde{\Lambda}_{cj} - \tilde{\Lambda}_{bi})^2 + ||\tilde{\Theta}_{cj} - \tilde{\Theta}_{bi}||^2}, \tag{7}$$

where $N_c$ is the number of scales in cluster $c$. The silhouette of grid cell $i$ in cluster $b$ compares its average distance to other grid cells within its own cluster against its average distance to its closest cluster:

$$\text{silhouette}_{bi} = \frac{\min\limits_{c \neq b} D^c_{bi} - D^b_{bi}}{\max\left[\min\limits_{c \neq b} D^c_{bi}, D^b_{bi}\right]}. \tag{8}$$

The denominator is a normalization factor that rescales the silhouette between $-1$ and 1. More positive values indicate better clustering. Out of the 200 clustering attempts, we select the one with largest average silhouette across all grid cells. Finally, we reject all clusters with 3 or fewer grid cells from further analysis. The remaining clusters are grid cell modules.

### VARYING VELOCITY GAIN MODEL

#### Simulation setup with a velocity gain gradient

These simulations use constant inhibition distance $l$ and a varying velocity gain $\alpha(z)$ (**Supp. Fig. 7b**). The functional form for $\alpha(z)$ is similar to that for $l(z)$ of the inhibition gradient model (see **Methods** of the main text), except it decreases with $z$ instead of increasing:

$$\alpha(z) = \left[\alpha_{\text{max}}^{\alpha_{\text{exp}}} + \left(\alpha_{\text{min}}^{\alpha_{\text{exp}}} - \alpha_{\text{max}}^{\alpha_{\text{exp}}}\right) \frac{z-1}{h-1}\right]^{1/\alpha_{\text{exp}}}, \tag{9}$$

which ranges from $\alpha_{\text{max}} = \alpha(1)$ to $\alpha_{\text{min}} = \alpha(h)$ with concavity tuned by $\alpha_{\text{exp}}$. More negative values of $\alpha_{\text{exp}}$ lead to greater concavity; for $\alpha_{\text{exp}} = 0$, we use the limiting expression $\alpha(z) = \alpha_{\text{max}}^{(h-z)/(h-1)} \alpha_{\text{min}}^{(z-1)/(h-1)}$.

Simulation initialization and time evolution proceed similarly to the inhibition gradient model, except we evolve 250 000 timesteps with a real rat trajectory before starting the main simulation phase, instead of 50 000 timesteps. Simulations with a velocity gain gradient tend to have transient configurations that persist longer before changing to a stable configuration, so a longer initialization period helps the main simulation start in a stable configuration.

#### Results with a velocity gain gradient

Simulations with a velocity gain gradient and excitatory coupling exhibit modularity, but grid scale and orientation relationships vary greatly among replicate simulations that use different random initial firing rates. Single neuron rate maps in **Supp. Fig. 7a** show that this model can produce a grid system with range of grid scales. Note that the population activity contains grids of the same scale for all networks because the inhibition distance is constant. Spatial scales are smaller at lower $z$ because they have higher velocity gain $\alpha$ (**Supp. Fig. 7b**) and translate their activity patterns in proportion to rat motion more rapidly. Plotting the spatial scales and orientations of all replicate simulations does not reveal strong clustering (**Supp. Fig. 7c**), but separate analysis of each replicate simulation allows us to identify mostly well-defined modules (**Supp. Fig. 7d**). However, scale ratios and orientation differences between adjacent modules do not cluster around preferred values (**Supp. Fig. 7e**).

To further investigate the dynamics of this model, we follow three pairs of adjacent networks in the **Supp. Video**, which corresponds to the replicate simulation shown in **Supp. Fig. 7a**. The movie depicts the activity overlays of these networks as the simulated rat explores its enclosure. We first consider the overlay between networks 1 and 2. Due to the "rigidity" provided by excitatory coupling as described in the main text, the population activities of these two networks remain in registry throughout the movie; thus, grid cells from these two networks have the same spatial scale and orientation and belong to the same module. Now consider the overlay between networks 3 and 4. For most of the movie, their population activities are in registry. However, the gradient in velocity gain prefers the more dorsal network (smaller $z$) to have an activity pattern that translates more rapidly. This effect can disrupt the rigidity imposed by coupling and, at $t \approx 470\,\mathrm{s}$, one pattern jumps along a lattice vector relative to the other. Such an anomaly implies that at least one of the two networks cannot have an activity pattern that translates proportionally to the entire rat trajectory, a requirement for faithful path-integration. Indeed, the spatial autocorrelation map for $z = 3$ in **Supp. Fig. 7a** shows a lack of grid-like symmetry. This example illustrates how a velocity gain gradient can disrupt grid cells in a way that an inhibition distance gradient does not—the latter does not resist the rigidity of excitatory coupling through different translation speeds of activity patterns.

Finally, consider the overlay between networks 7 and 8 in the **Supp. Video**. Here, the two activity patterns remain rotated relative to each other, with little registry. Coupling causes the activity peaks of network 8 to preferentially excite the corresponding areas of network 7, but since there are few peaks in those areas, the effect of coupling is weak between these two networks. Therefore, the activity patterns can freely glide relative to each other, each translating proportionally with animal motion but with different speeds preferred by different velocity gains. Indeed, single neuron spatial rate maps for $z = 7$ and 8 show different scales (**Supp. Fig. 7a**), which identifies this lack of registry as a mechanism for producing interfaces between grid modules. However, this mechanism does not enforce how quickly one pattern glides relative to the other and thus does not lead to preferred scale ratios (**Supp. Fig. 7e**). It does require that activity patterns stay rotated relative to each other, which may explain the abundance of large orientation differences $>15^\circ$ between modules (**Supp. Fig. 7e**).

Thus, excitatory coupling with a velocity gain gradient can produce grid modules, but, in contrast to the model with varying inhibition distance, the velocity gain gradient model does not favor certain scale ratios and orientation differences.

### SUPPLEMENTARY VIDEO CAPTION

Dynamic relationships between adjacent networks in the model with varying velocity gain $\alpha(z)$. The movie spans the last 100 s of the simulation displayed in **Supp. Fig. 7a**. Top left: accumulated rat trajectory (gray curve) with current rat position (blue dot). Top right, bottom left, and bottom right: activity overlays between adjacent networks with the network at smaller (larger) $z$ depicted in magenta (green), so white indicates regions of activity in both networks. White scale bar, 50 neurons. Black scale bar, 50 cm.

### SUPPLEMENTARY FIGURES

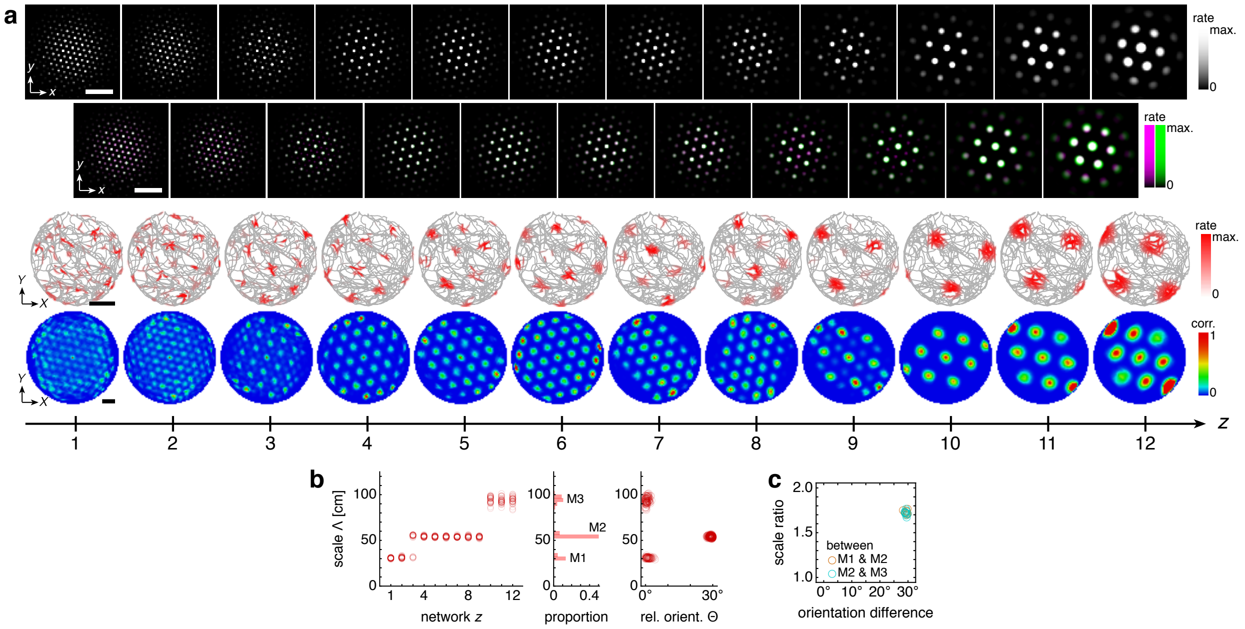


**Supplementary Figure 1**. Model with coupling spread $d(z)$ varying proportionally to inhibition distance $l(z)$ produces modules with the same scale ratios and orientation differences as the standard model with fixed $d$. We choose $d(z) = l(z)$ with all other simulation parameters unchanged (**Table 1** of the main text). (**a**) Representative simulation. Top row: network activities at the end of the simulation. Second row: activity overlays between adjacent networks depicted in the top row. In each panel, the network at smaller (larger) $z$ is depicted in magenta (green), so white indicates regions of activity in both networks. Third row: spatial rate map of a single neuron for each $z$ superimposed on the animal's trajectory. Bottom row: spatial autocorrelations of the rate maps depicted in the third row. (**b,c**) Statistics based on 10 replicate simulations subject to a gridness cutoff of 0.6. (**b**) Left: spatial grid scales $\Lambda(z)$. Middle: histogram for $\Lambda$ collected across all depths $z$. Grid cells are clustered into three modules. Right: spatial grid orientations $\Theta$ relative to the grid cell in the same simulation with largest scale. (**c**) Spatial grid scale ratios and orientation differences between adjacent modules. White scale bars, 50 neurons. Black scale bars, 50 cm.

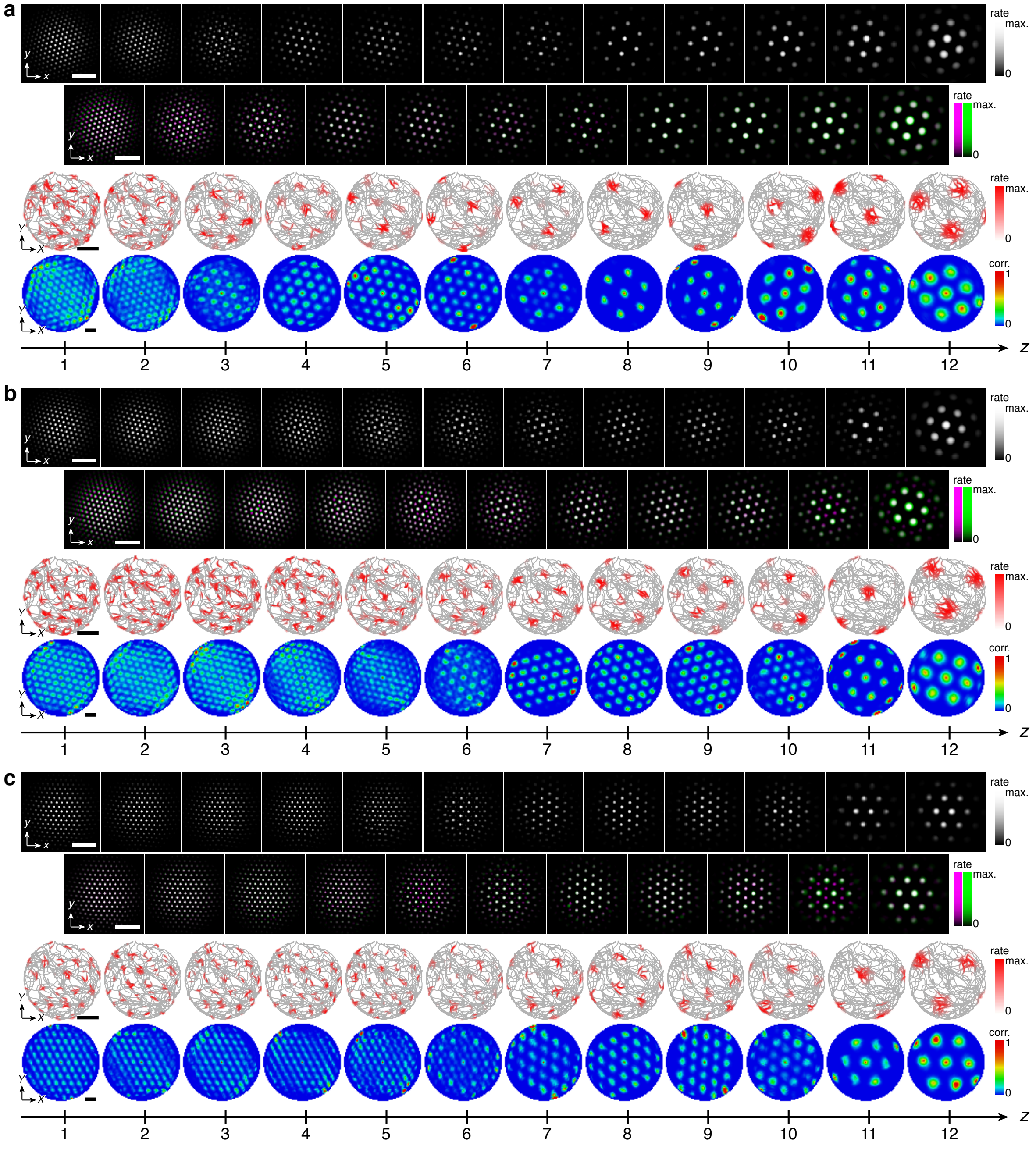


**Supplementary Figure 2**. Representative network activities and single neuron rate maps corresponding to **Fig. 3** of the main text. (**a**) Inhibition distance scaling exponent $l_{\mathrm{exp}} = 0$, corresponding to **Fig. 3a** of the main text. Top row: network activities at the end of the simulation. Second row: activity overlays between adjacent networks depicted in the top row. In each panel, the network at smaller (larger) $z$ is depicted in magenta (green), so white indicates regions of activity in both networks. Third row: spatial rate map of a single neuron for each $z$ superimposed on the animal's trajectory. Bottom row: spatial autocorrelations of the rate maps depicted in the third row. (**b**) Same as **a**, but for $l_{\mathrm{exp}} = -2$, corresponding to **Fig. 3b** of the main text. (**c**) Same as **a**, but for bidirectional point-to-point coupling with spread $d = 1$ and strength $u_{\mathrm{mag}} = 0.4$, corresponding to **Fig. 3c** of the main text. Other parameter values are in **Table 1** of the main text. White scale bars, 50 neurons. Black scale bars, 50 cm.

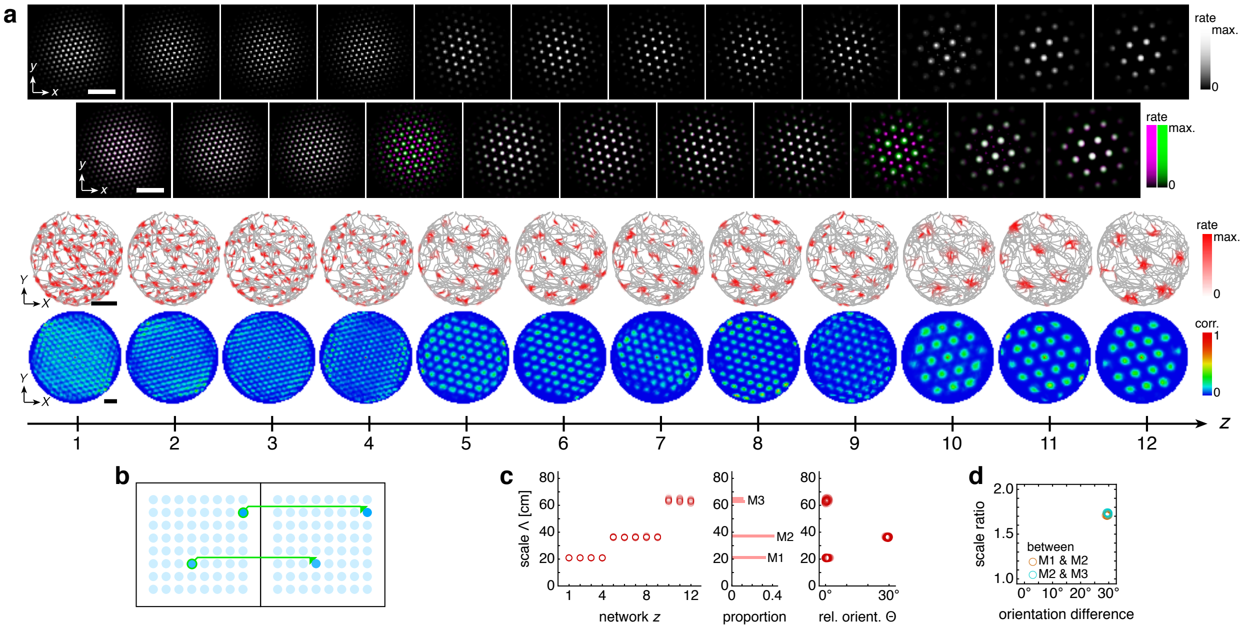


**Supplementary Figure 3**. Model with dorsal-to-ventral point-to-point coupling produces modules with the same scale ratios and orientation differences as the standard model. (**a**) Representative simulation. Top row: network activities at the end of the simulation. Second row: activity overlays between adjacent networks depicted in the top row. In each panel, the network at smaller (larger) $z$ is depicted in magenta (green), so white indicates regions of activity in both networks. Third row: spatial rate map of a single neuron for each $z$ superimposed on the animal's trajectory. Bottom row: spatial autocorrelations of the rate maps depicted in the third row. (**b**) Schematic of dorsal-to-ventral point-to-point coupling. The neuron at position $(x, y)$ in network $z$ excites only the neuron at $(x, y)$ in network $z + 1$. (**c,d**) Statistics based on 10 replicate simulations subject to a gridness cutoff of 0.6. (**c**) Left: spatial grid scales $\Lambda(z)$. Middle: histogram for $\Lambda$ collected across all depths $z$. Grid cells are clustered into three modules. Right: spatial grid orientations $\Theta$ relative to the grid cell in the same simulation with largest scale. (**d**) Spatial grid scale ratios and orientation differences between adjacent modules. Coupling spread $d = 1$ and coupling strength $u_{\mathrm{mag}} = 0.8$. Other parameter values are in **Table 1** of the main text. White scale bars, 50 neurons. Black scale bars, 50 cm.

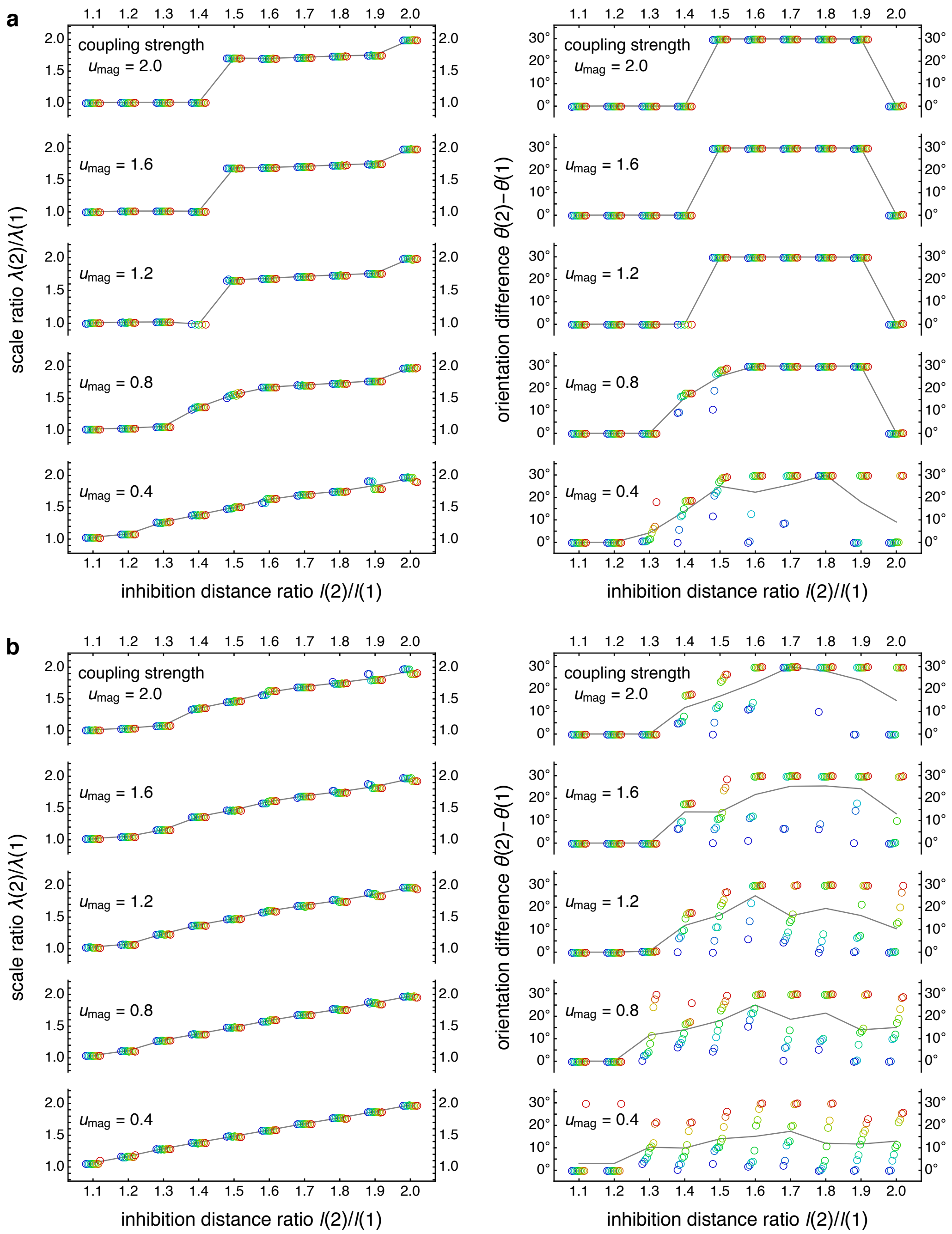


**Supplementary Figure 4**. Raw scale ratio and orientation difference data used to produce the phase diagrams of **Fig. 4** of the main text. Simulations have only 2 network depths. We systematically change coupling strength $u_{\mathrm{mag}}$ and inhibition distance ratio $l(2)/l(1)$ while fixing the mean of $l(1)$ and $l(2)$ to be 9. Network size is $n \times n = 230 \times 230$. (**a**) Data for coupling spread $d = 6$. Left: network spacing ratios $\lambda(2)/\lambda(1)$; Right: network orientation differences $\theta(2) - \theta(1)$. For each $l(2)/l(1)$ and $u_{\mathrm{mag}}$, 10 replicate simulations subject to a gridness cutoff of 0.6 are represented by circles with colors corresponding across the two panels. Small horizontal offsets are introduced for clarity, and each set of replicate simulations is ordered by $\theta(2) - \theta(1)$. Gray lines track mean values as a function of $l(2)/l(1)$. (**b**) Same as **a**, but for coupling spread $d = 12$. Parameter values are in the **Simulation setup** section and in **Table 1** of the main text.

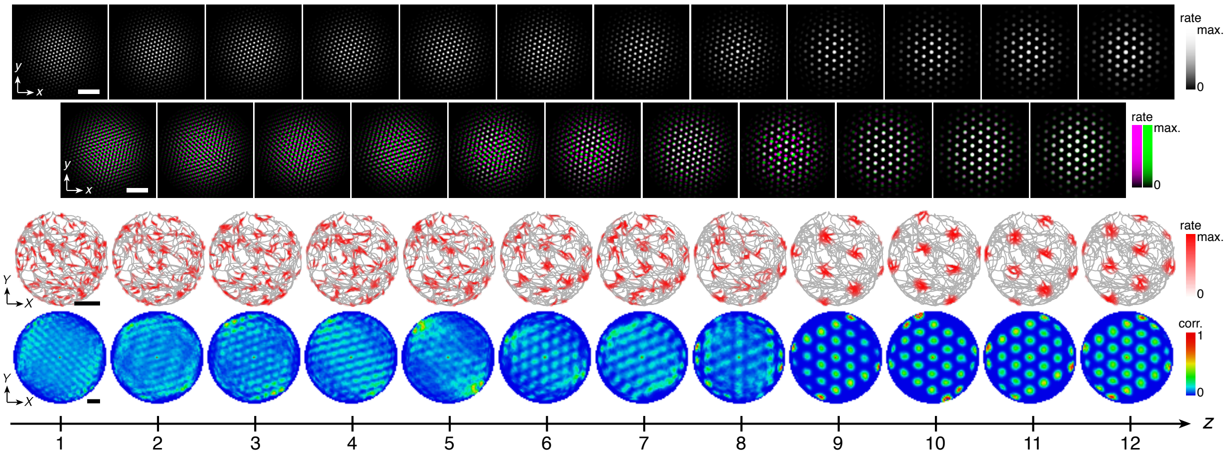


**Supplementary Figure 5**. (**a**) Representative network activities and single neuron rate maps for the discommensurate system, corresponding to **Fig. 5** of the main text. Top row: network activities at the end of the simulation. Second row: activity overlays between adjacent networks depicted in the top row. In each panel, the network at smaller (larger) $z$ is depicted in magenta (green), so white indicates regions of activity in both networks. Third row: spatial rate map of a single neuron for each $z$ superimposed on the animal's trajectory. Bottom row: spatial autocorrelations of the rate maps depicted in the third row. White scale bars, 50 neurons. Black scale bars, 50 cm. (**b**) Clustering of spatial scales and orientations for all replicate simulations. Due to 6-fold lattice symmetry, orientation is a periodic variable modulo 60°. Different colors indicate separate modules. Simulations use network size 230 × 230, maximum inhibition distance $l_{\max} = 10$, and coupling spread $d = 12$.

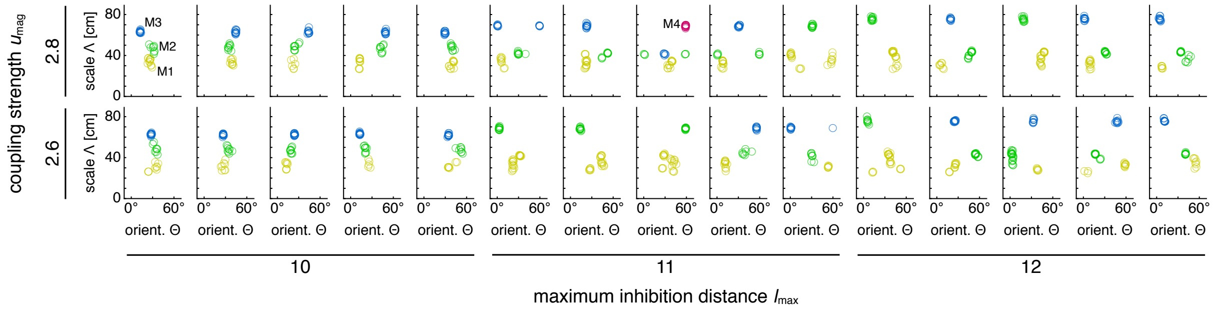


**Supplementary Figure 6**. Clustering of scales and orientations for simulations spanning a range of parameters, corresponding to **Fig. 6** of the main text. Due to 6-fold lattice symmetry, orientation is a periodic variable modulo 60°. Different colors indicate separate modules. Simulations use network size 230 × 230 and coupling spread $d = 12$.

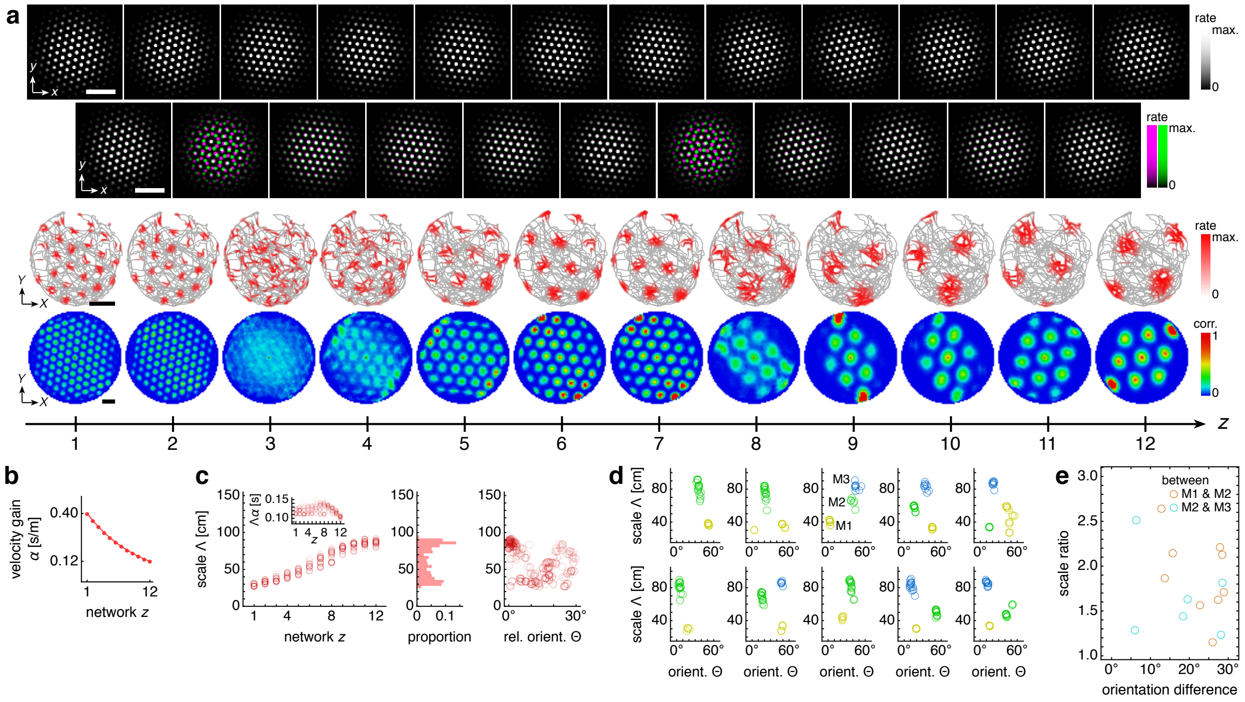

**Supplementary Figure 7**. Simulations with a varying velocity gain $\alpha(z)$ and constant inhibition distance $l$ produce modules that do not exhibit preferred relationships. (**a**) Representative simulation. Top row: network activities at the end of the simulation. Second row: activity overlays between adjacent networks depicted in the top row. In each panel, the network at smaller (larger) $z$ is depicted in magenta (green), so white indicates regions of activity in both networks. Third row: spatial rate map of a single neuron for each $z$ superimposed on the animal's trajectory. Bottom row: spatial autocorrelations of the rate maps depicted in the third row. (**b**) Velocity gain profile $\alpha(z)$. (**c**–**e**) Data from 10 replicate simulations. (**c**) Left: spatial grid scales $\Lambda(z)$. For each network, there are up to 30 red circles corresponding to 3 neurons recorded during each simulation. Inset: $\Lambda(z)$ multiplied by the velocity gain $\alpha(z)$. Middle: histogram for $\Lambda$ collected across all networks. Right: spatial grid orientations $\Theta$ relative to the grid cell in the same simulation with largest scale. (**d**) Distribution of spatial grid scales and orientations relative to an arbitrary axis for each replicate simulation. Due to hexagonal symmetry, orientation is a periodic variable modulo 60°. Different colors indicate separate modules. The ninth panel corresponds to the overlays in **a**. (**e**) Spatial grid scale ratios and orientation differences between adjacent modules. Maximum velocity gain $\alpha_{\text{max}} = 0.40\,\text{s/m}$, minimum velocity gain $\alpha_{\text{min}} = 0.12\,\text{s/m}$, and scaling exponent $\alpha_{\text{exp}} = 0$. Network size $n \times n = 174 \times 174$, coupling spread $d = 12$, coupling strength $u_{\text{mag}} = 0.6$, and inhibition distance $l = 6$. Other parameter values are in **Table 1** of the main text. White scale bars, 50 neurons. Black scale bars, 50 cm.

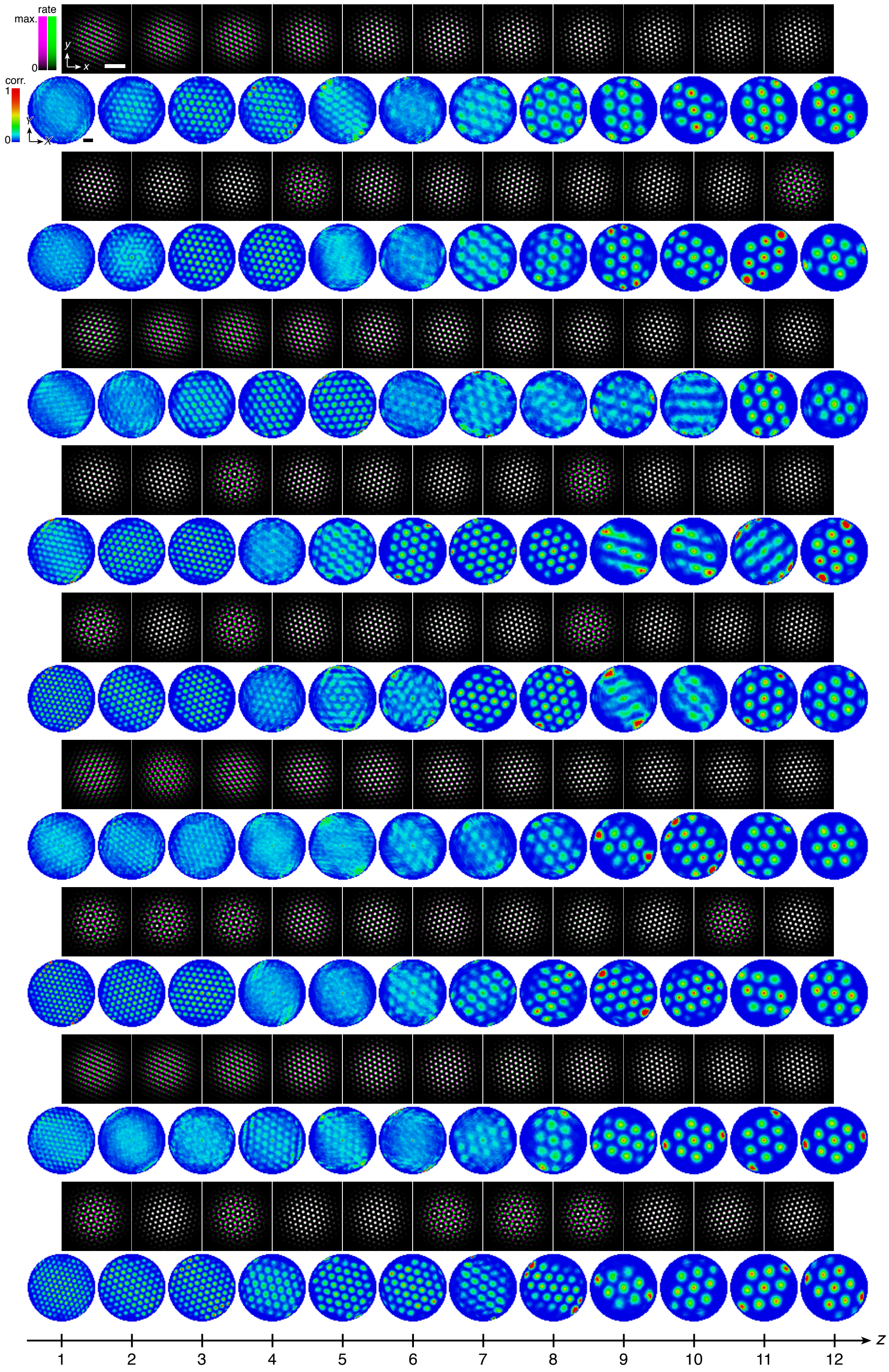


**Supplementary Figure 8**. Activities of replicate simulations with a varying velocity gain contributing to **Supp. Fig. 7**, except for the one shown in **Supp. Fig. 7a**. Top row in each panel: activity overlays between adjacent networks with the network at smaller (larger) $z$ depicted in magenta (green). Bottom row in each panel: autocorrelations of spatial rate maps. White scale bar, 50 neurons. Black scale bar, 50 cm.

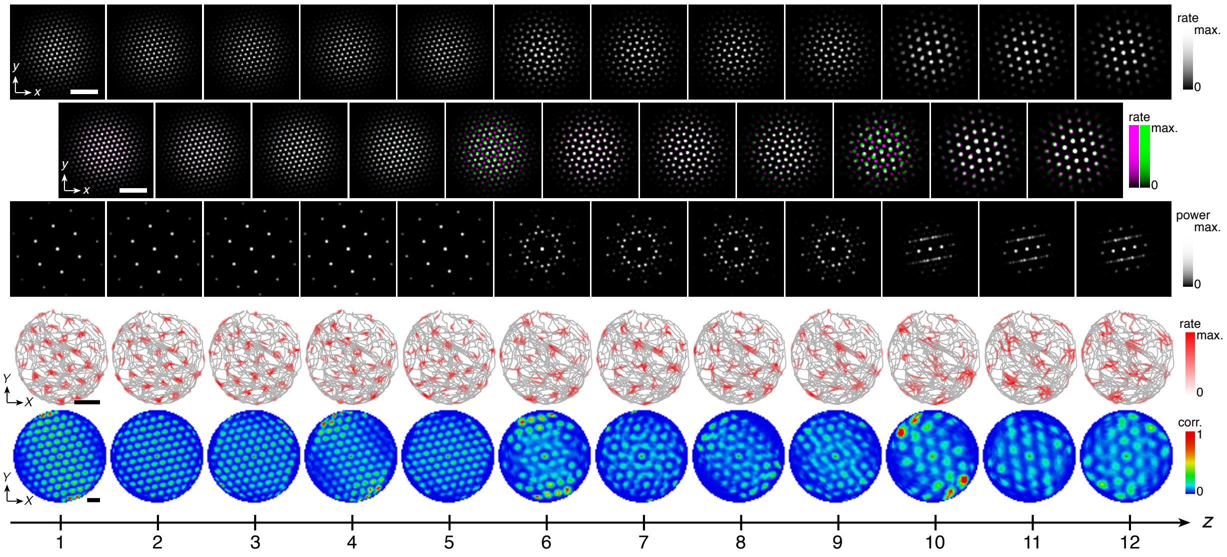


**Supplementary Figure 9**. Quasicrystal approximant grids in a representative simulation with dorsal-to-ventral coupling. Top row: network activities at the end of the simulation. From $z = 6$ to 9, activity peaks form the vertices of a square-triangle tiling that is a dodecagonal quasicrystal approximant [6, 7]. This tiling is labeled $(3^6; 3^2.4.3.4)$ based on the type and order of regular polygons that meet at its vertices [8]. From $z = 10$ to 12, the activity patterns demonstrate 2-fold dihedral symmetry. Second row: activity overlays between adjacent networks depicted in the top row. In each panel, the network at smaller (larger) $z$ is depicted in magenta (green), so white indicates regions of activity in both networks. Third row: Fourier power spectra for network activities with the origin at the center of each image and the edges cropped. Note that the $z = 8$ and 9 spectra approach 12-fold symmetry, as expected from a dodecahedral quasicrystal approximant. Fourth row: spatial rate map of a single neuron for each $z$ superimposed on the animal's trajectory. Bottom row: spatial autocorrelations of the rate maps depicted in the third row. Network size $n \times n = 230 \times 230$, coupling spread $d = 2$, coupling strength $u_{\mathrm{mag}} = 0.6$, maximum inhibition distance $l_{\mathrm{max}} = 10$, and velocity gain $\alpha = 0.2\,\mathrm{s/m}$. Other parameter values are in **Table 1** of the main text. White scale bars, 50 neurons. Black scale bars, 50 cm.
